# Human motor cortical gamma activity relates to GABAergic signalling and to behaviour

**DOI:** 10.1101/2021.06.16.448658

**Authors:** Catharina Zich, Magdalena Nowak, Emily L Hinson, Andrew J Quinn, Mark W Woolrich, Charlotte J Stagg

## Abstract

Gamma activity (γ, >30 Hz) is universally demonstrated across brain regions and species. However, the physiological basis and functional role of γ sub-bands (slow-γ, mid-γ, fast-γ) have been predominantly studied in rodent hippocampus; γ activity in the human neocortex is much less well understood.

Here we combined neuroimaging and non-invasive brain stimulation to examine the properties of γ activity sub-bands in the primary motor cortex (M1), and their relationship to both local GABAergic activity and to motor learning. In 33 healthy individuals, we quantified movement-related γ activity in M1 using magnetoencephalography, assessed GABAergic signaling using transcranial magnetic stimulation (TMS), and estimated motor learning via a serial reaction time task.

We characterised two distinct γ sub-bands (slow-γ, mid-γ) which show movement-related increase in activity during unilateral index finger movements and are characterised by distinct temporal-spectral-spatial profiles. Bayesian correlation analysis revealed strong evidence for a positive relationship between slow-γ (∼30-60Hz) peak frequency and endogenous GABA signalling during movement preparation (as assessed using the TMS-metric short interval intracortical inhibition). There was also moderate evidence for a relationship between power of the movement-related mid-γ activity (60-90Hz) and motor learning. These relationships were neurochemically- and frequency-specific.

These data provide new insights into the neurophysiological basis and functional roles of γ activity in human M1 and allow the development of a new theoretical framework for γ activity in the human neocortex.

**Significance Statement:** Gamma (γ) activity is ubiquitous in the brain, yet our understanding of the mechanisms and function of γ activity in the human neocortex, and particularly in the human motor cortex, is limited. Using a multimodal approach, we characterised two patterns of movement-related γ activity in the human motor cortex (slow-γ and mid-γ), with different spatial, temporal and spectral properties. Slow-γ peak frequency was correlated to local GABA-A activity, whereas mid-gamma power predicted performance in a subsequent motor learning task. Based on these findings and previous research, we propose a theoretical framework to explain how human motor cortical γ activities may arise and their potential role in plasticity and motor learning, providing new hypotheses to be tested in future studies.

**Key Points:**

- We combined neuroimaging (i.e. MEG) and non-invasive brain stimulation (i.e. TMS) to examine the properties of γ activity sub-bands in the primary motor cortex.
- Two distinct γ sub-bands (slow-γ, mid-γ) show a movement-related increase in activity during finger movements and are characterised by distinct temporal-spectral-spatial profiles.
- We found strong evidence for a positive relationship between slow-γ (∼30-60Hz) peak frequency and endogenous GABA signalling during movement preparation (as assessed using the TMS-metric short interval intracortical inhibition).

## 1 Introduction

Activity within the gamma band (γ, >30 Hz) is ubiquitous across the mammalian brain. This broad frequency band is commonly divided into three sub-bands: slow-γ (human: ∼30-60Hz: rodent: ∼30-50Hz), mid-γ (human: ∼60-90Hz; rodent: ∼50-100 Hz) and fast-γ (human: >90 Hz; rodent: >100 Hz). To date, γ activity has been mainly explored in rodent hippocampus, where the sub-bands of this activity arise from separate locations (Schomburg et al., 2014; Lasztóczi and Klausberger, 2016), reflect distinct microcircuits (Bragin et al., 1995; Colgin et al., 2009; Fernández-Ruiz et al., 2017), and have different functional roles (Carr and Frank, 2012; Colgin, 2015). Considerably less is known about the neurophysiological bases and functional roles of the γ sub-bands in the human neocortex.

In the human motor system, a movement-related increase in γ power (γ event-related synchronization [γ ERS]) has been described (Crone et al., 1998; Pfurtscheller and Lopes da Silva, 1999; Canolty et al., 2006; Cheyne et al., 2008; Muthukumaraswamy, 2010, 2011; Cheyne, 2013). This has been most frequently reported for the mid-γ band (i.e. mid-γ ERS), which occurs only during actual rather than imagined movement (Muthukumaraswamy, 2011), shows spatial specificity to the primary motor cortex (M1, (Crone et al., 2006)) and temporal specificity to the time of the movement. Mid-γ has been suggested to play a role in afferent proprioceptive feedback or relate to more active motor control processes (Miller et al., 2010; Muthukumaraswamy, 2010), and its pro-kinetic role has been demonstrated in a number of studies (Joundi et al., 2012; Swann et al., 2016, 2018). In humans, slow-γ has been considerably less well characterised. Only few studies have reported movement-related slow-γ activity in the M1 (Crone et al., 1998; Szurhaj et al., 2006). Their data suggested that slow-γ has a distinct spatio-temporal profile and plays a functional role in synchronising activity of neuronal populations involved in movement.

Slow-γ and mid-γ likely share some physiological similarities. Empirical animal studies and computational modelling have demonstrated that GABAergic interneuron-mediated inhibition of pyramidal cell activity generates γ activity in M1 (Gonzalez-Burgos and Lewis, 2008; Sohal et al., 2009; Buzsáki and Wang, 2012). Despite initial findings in humans relating γ activity in M1 to GABA (Gaetz et al., 2011) it remains to be determined how closely these data, which arise from in vitro and invasive in vivo recordings, translate into the γ ERS seen in human electrophysiological recordings. However, given the wealth of data implicating changes in M1 GABAergic activity during motor learning (Stagg et al., 2011), y activity may reflect a mechanism by which decreases in local GABAergic signalling mediates behavioural improvements. In line with this hypothesis, our group recently demonstrated that 75 Hz tACS applied to M1 leads to a significant reduction in local GABA-A activity, as assessed by transcranial magnetic stimulation (TMS, (Nowak et al., 2017)). Moreover, this 75 Hz tACS-induced change in GABA-A activity was closely correlated with an individual’s motor learning ability.

The present study aims to investigate the relationship between M1 GABAergic signalling, M1 γ activity and motor learning in humans. We characterized M1 γ activity in 33 young healthy individuals during unilateral index finger movement using magnetoencephalography (MEG), to address the hypotheses that an individual’s M1 movement-related γ activity is related to TMS measure of local GABA-A activity and the ability to learn a motor task.

## 2 Methods

### 2.1 Participants

33 individuals (age 24.9 years, range: 21–30 years; 14 male) gave their informed consent to participate in the study in accordance with Central University Research Ethics Committee approval (University of Oxford; MSD-IDREC-C2–2014-026 and MSD-IDREC-C1-2015-010). All participants were right-handed as assessed by the Edinburgh Handedness Inventory (Oldfield, 1971), had no history of neurological or psychiatric disorders, no metal implants, and reported no other contraindications to TMS or MEG. A subset of these data formed part of a previous publication (Nowak et al., 2017).

### 2.2 Experimental Design

All individuals completed all parts of the study on the same day in the following order: MEG data recording during rest and during a Go/NoGo task, a motor learning task (ML) and a response time task (RT) outside the MEG, and TMS during rest and RT task (**Fig. 1a**).

**Fig. 1.**
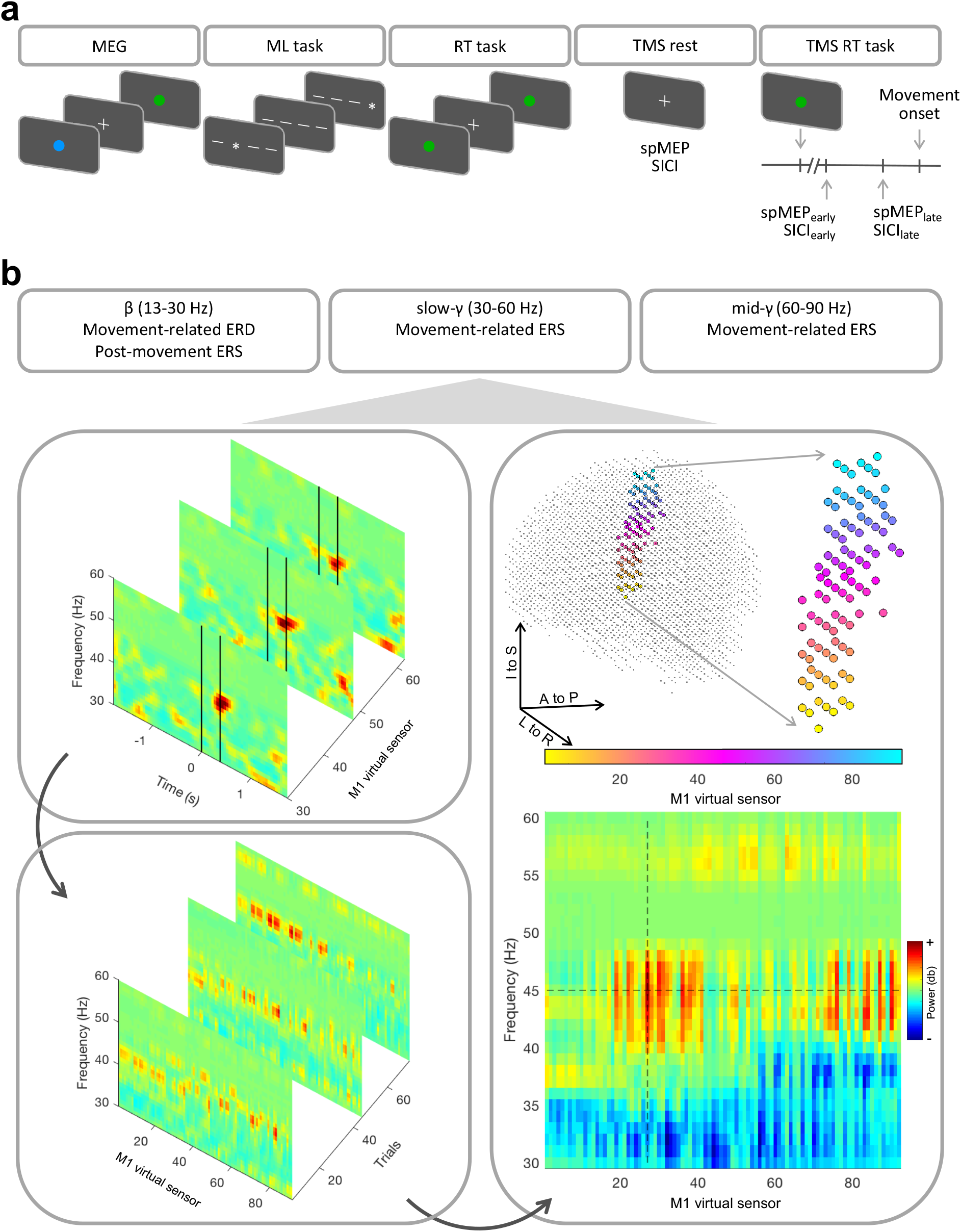
(a) Experimental timeline. MEG data were recorded while individuals performed a Go/NoGo task (70 trials). Each trial started with a preparation cue (blue circle, for 200 ms), followed by a cue-stimulus interval (fixation cross, 1000 ms) and the Go (green circle, 80% of the trials) or NoGo (red circle) cue. After the MEG participants performed a visually cued motor sequence learning (ML) task (4 fingers, 10-item sequence, 3 repeats per block, 15 blocks). Individuals also performed a simple response time (RT) task (20 trials) to determine their individual RT. Each trial started with a green circle indicating to perform an index finger abduction followed by a fixation cross indicating the inter-trial interval. TMS measures (single-pulse MEP [spMEP] and short interval intracortical inhibition [SICI]) were performed at rest and during the RT task. During the RT task, spMEP and SICI were obtained at two different timings during movement preparation: an early time point (25% of mean RT, spMEP_early_, SICI_early_) and a late time point (65% of mean RT, spMEP_late_, SICI_late_). (**b**) Pipeline used to identify the individuals’ peak frequency for β ERD, β ERS, slow-γ ERS, mid-γ ERS, exemplary for one individual for slow-γ ERS. For each individual trial the power from movement onset to movement offset (movement offset to movement offset + 1 s for β ERS), as determined by EMG and indicated by the black vertical lines, was averaged for each frequency band and virtual sensors within the M1 separately (top left). This resulted in movement-related power (post-movement-related power for β ERS) for each frequency, M1 virtual sensor and trial (bottom left). The maximum (minimum for β ERD) of the average across trials (right) indicates the individuals’ power, peak frequency (dashed horizontal line) and virtual sensor within M1 (dashed vertical line). Virtual sensors are ordered based on MNI z-coordinates and color-coded accordingly (e.g. small index = yellow = inferior).

#### MEG data acquisition

MEG data were acquired with a whole-head 306-channel Elekta Neuromag system (204 planar gradiometers, 102 magnetometers). Concurrent surface electromyography (EMG) of the right extensor digitorum communis and first dorsal interosseous (FDI) muscle were recorded using bipolar surface electrodes. Both MEG and EMG data were sampled at 1000 Hz with a band-pass filter of 0.03-330 Hz. Head position was continuously monitored with respect to the MEG sensors using four head-position (HPI) coils. The locations of HPI coils and of three anatomical fiducials (the nasion and two preauricular points) were digitized using a 3D tracking system (Polhemus, EastTrach 3D) to define the subject-specific cartesian head coordinate system. In addition, vertical and horizontal electrooculogram electrodes were used to allow for detection and removal of eye-blink artefacts. MEG data were acquired during a Go/NoGo task. A blue circle cue, presented for 200 ms, instructed participants to prepare for an abduction of the index finger of their right hand. The cue was then replaced by a fixation cross for 1000 ms (cue - stimulus interval). A subsequent visual stimulus presented for 200 ms (coloured circle: green for Go or red for NoGo) indicated whether they should perform (Go) or withhold (NoGo) the prepared response. Participants were instructed to respond as quickly as possible on the Go trials. The stimulus was then replaced by a fixation cross for a duration that varied randomly between 2000 and 4000 ms (inter-trial interval). The task consisted of a total of 70 trials and lasted ∼5 min. NoGo trials (20% of all trials) were introduced to encourage participants’ attention to the task. Stimuli were generated using the MATLAB Psychophysics Toolbox version 3.0 package (Brainard, 1997) and back-projected (Panasonic DLP Projector, PT D7700E) onto a screen at a viewing distance of 120 cm with a spatial resolution of 1024 by 768 pixels and a refresh rate of 60 Hz.

#### Motor learning task

Outside the MEG, individuals performed a visually cued motor sequence learning task which has been described previously (Nowak et al., 2017). Briefly, individuals were presented with four horizontal bars on a screen, each of which corresponded to a key on the keyboard. When a bar changed into an asterisk, individuals were instructed to press the corresponding key as quickly and accurately as possible. The task included sequence blocks consisting of three repeats of a 10-item sequence. The first and 15th blocks consisted of 30 visual cues presented in a random order.

#### Response time task

Individuals performed also a simple RT task consisting of 20 trials to characterize their individual RT for the subsequent TMS measurement. Individuals were instructed to respond to a visual Go signal (coloured green circle) by performing an index finger abduction of the right hand as quickly as possible. Visual stimuli appeared at random intervals (5-7 s) and the individuals were instructed to avoid anticipation of the Go signal and to relax their hand while the fixation cross was displayed on the screen. Stimuli were generated using the MATLAB Psychophysics Toolbox version 3.0 package (Brainard, 1997). Surface EMG was recorded via disposable neonatal ECG electrodes (Henley’s Medical) from the FDI of the right hand using a belly-tendon montage with a ground electrode over the ulnar styloid process. Signals were sampled at 5 kHz, amplified, filtered (10-1000 Hz), and recorded using a CED 1902 amplifier, a CED micro1401 A/D converter, and Signal software version 3.13 (Cambridge Electronic Design).

#### TMS data acquisition

All TMS data were acquired using a monophasic BiStim machine connected to a 70 mm figure-of-eight coil (Magstim). The left M1 FDI motor hotspot, i.e. the position where single-pulse motor evoked potentials (spMEPs) could be elicited in the in the right FDI muscle at the lowest stimulator intensity, was targeted. The TMS coil was held at 45° to the midsagittal line with the handle pointing posteriorly. The hotspot was marked on a tight-fitting cap to ensure reproducible coil positioning. Surface EMG data were recorded in the same manner as during the RT task.

First, resting motor threshold (rMT) and active motor threshold (aMT) were determined. rMT was defined as the minimum stimulus intensity required for eliciting spMEPs of ∼1 mV peak-to-peak amplitude in at least 5/10 trials in the relaxed FDI muscle. aMT was defined as the minimum stimulus intensity necessary to evoke spMEPs of ∼200 μV peak-to-peak amplitude in at least 5/10 trials while individuals maintained ∼30% of the maximum contraction of the FDI.

Local GABA-A synaptic activity was assessed using short interval intracortical inhibition (SICI) with an interstimulus interval of 2.5 ms (Kujirai et al., 1993; Di Lazzaro et al., 2005). The conditioning stimulus was set at 70% of aMT and the test stimulus at rMT. spMEP and SICI were measured in pseudorandomized order with fifteen trials per condition, both at rest and during the pre-movement period of the simple RT task (same RT task as performed prior to TMS). TMS measures were collected at two different times during movement preparation: an early time point (25% of mean RT) and a late time point (65% of mean RT), resulting in four different pre-movement protocols: spMEP_early_, spMEP_late_, SICI_early_ and SICI_late_. The 25% and 65% RT were adjusted to each individual’s mean RT according to a previously described procedure (Murase et al., 2004; Hummel et al., 2009). Fifteen trials per condition and time point were recorded.

### 2.3 Data analysis

#### MEG data analysis

External noise was reduced from MEG data by means of spatio-temporal signal-space separation (TSSS) and head movements (detected using HPI coils) corrected, both using MaxMove software as implemented in MaxFilter version 2.1 (Elekta Neromag, Elekta, Stockholm, Sweden). Further MEG data analyses were performed using the in-house OHBA Software Library (OSL: https://ohba-analysis.github.io/osl-docs/) version 2.2.0. Registration between a structural MRI template, i.e. MNI152 standard-space T1-weighted average structural template image, and MEG data was performed with RHINO (Registration of Headshapes Including Nose in OSL) using nose and fiducial landmarks for coregistration and a single shell as forward model.

Continuous data were down-sampled to 500 Hz. Further, a band-pass filter (5-245 Hz) and several notch filters were applied (49-55 Hz, 99-101 Hz, 149-151 Hz, 199-201 Hz). A wider notch filter around 50 Hz was used to supress 50 Hz line noise and a 53 Hz artefact present in this dataset caused by the HPI coils. Time segments containing artefacts were identified by using generalized extreme studentized deviate method (GESD (Rosner, 1983)) at a significance level of 0.05 with a maximum number of outliers limited to 20% of the data on the standard deviation of the signal across all sensors in 1 s non-overlapping windows. The windows corresponding to outliers were excluded from all further analysis. Further denoising was applied using independent component analysis (ICA) using temporal FastICA across sensors (Hyvarinen, 1999). 62 independent components were estimated and components representing stereotypical artefacts such as eye blinks, eye movements, and electrical heartbeat activity were manually identified and regressed out of the data. Data then were filtered into three frequency bands (β 13-30 Hz, slow-γ 30-60 Hz, mid-γ 60-90 Hz) and the following processing steps were performed separately for the three frequency bands.

Magnetometers and Planar-Gradiometers were normalized by computing the eigenvalue decomposition across sensors within each coil type and dividing the data by the smallest eigenvalue within each (Woolrich et al., 2011). Data were projected onto an 8 mm grid in source space (resulting in 3559 virtual sensors) using a Linearly Constrained Minimum Variance (LCMV) vector beamformer (Van Veen and Buckley, 1988; Woolrich et al., 2011). Beamformer weights were estimated across an 8 mm grid cast within the inner-skull of the MNI152 brain. A covariance matrix was computed across the whole time-course and was regularized to 50 dimensions using principal component analysis (PCA) rank reduction (Quinn et al., 2018).

Epochs were defined relative to the movement onset (movement offset for β ERS) as identified by surface EMG. To identify movement onset and offset EMG data were first high-pass filtered at 10 Hz. EMG data were then segmented from -1 s to 3 s relative to the Go stimuli, and the envelope (root mean square, window = 80 ms), computed. Using a non-overlapping moving standard deviation (movement onset: window = 24 ms, direction = forward; movement offset: window = 120 ms, direction = backwards) movement onset and offset were defined as the first window exceeding the threshold (three standard deviations of the EMG activity between -600 ms to -200 ms relative to Go stimuli). Trials were excluded when the envelope, the reaction time (i.e. time between Go stimuli and movement onset), or the movement time (i.e. time between movement onset and offset) were identified as outliers using GESD at a significance level of 0.05. This approach results in 45.59 (SD = 4.88) out of 56 epochs per individual. MEG data were segmented from -2 s to 2 s relative to movement onset (movement offset for β ERD).

Time-frequency analysis was applied to single trials and virtual sensors using dpss-based multitaper (window = 1.6 s, steps = 200 ms) with a frequency resolution of 1 Hz. Segments were baseline corrected (−1 s to -0.5 s, [-1.5 s to -1 s for β ERD]) using the mean baseline across all trials. This procedure results in trial-by-trial time-frequency decomposition for each of the 3559 virtual sensors. To detect the individual peak frequency within each frequency band only the virtual sensors within M1, following the Desikan-Killiany atlas, (N = 92) were considered. Trial-wise movement-related power was obtained by averaging across time, i.e. from movement onset to movement offset (from movement offset to movement offset + 1 s for β ERS) and then averaged across trials. The maximum (minimum for β ERD) of the resulting two-dimensional matrix containing the power between movement on- and offset across trials at each frequency within the frequency band at each virtual sensor was defined as an individual’s power and defined the individual’s peak frequency and virtual sensor (**Fig. 1b**).

To illustrate the spatial properties of movement-related responses we computed the movement-related power (as above, first averaging across time, i.e. from movement onset to movement offset [from movement offset to movement offset + 1 s for β ERS] and then averaged across trials) at the individual’s peak frequency for each of the 3559 virtual sensors.

#### Motor learning data analysis

To derive an accurate motor learning score, individual RTs (i.e. time from cue onset to the correct button press) were first evaluated. Anticipatory responses (i.e. those that occurred before the cue) and outliers (i.e. RTs outside of the mean value ± 2 SD per each block) were discarded. A motor learning score was calculated for each individual as a percentage change from the RT in the first sequence block (block 2) to blocks 10-14, when the learning plateaued (Stagg et al., 2011). Thus, more negative motor learning scores represent better performance. One individual was excluded from this analysis due to noncompliance with instructions.

#### Response time task data analysis

EMG data were analysed online using Signal software version 3.13 (Cambridge Electronic Design). RT was identified for each trial as the time interval between the Go signal and the onset of EMG activity recorded above the FDI muscle (i.e. mV first data point where EMG amplitude >0.1 mV). The mean RT was then used in the remainder of that experimental session to calculate the timing of the pre-movement TMS pulses.

#### MEP data analysis

Trials were excluded if the test pulse alone failed to elicit a reliable MEP (amplitude <0.1 mV), there was precontraction in the target FDI muscle (EMG amplitude >0.1 mV in the 80 ms preceding the pulse), or, for the pre-movement TMS measures, EMG onset coincided with TMS pulse or no response was made. The peak-to-peak amplitude for each MEP was then calculated. Any MEPs outside of the mean value ± 2 SD for each condition for each block were excluded. Next, a single iteration of Grubbs’ test with a significance level of 0.05 was performed for each TMS condition separately and any significant outliers excluded. Collectively, these rejection criteria resulted in the exclusion of <5 trials per individual in any condition. SICI and ICF were expressed as a ratio of the mean conditioned MEP amplitude to the mean unconditioned MEP amplitude. For the pre-movement data, the TMS measures were analysed separately for each pre-movement time point (25% and 65% RT).

### 2.4 Statistical analysis

All Frequentist statistics were conducted as implemented in SPSS version 25 (SPSS Inc, Chicago, IL, USA). As Bayesian inference allows multiple hypotheses to be tested and can calculate the probability that one hypothesis is true relative to another hypothesis, correlation analysis was performed using Bayesian inference (JASP, JASP Team 2019, version 0.11.1) with default priors after outlier removal. The Bayes factors (BFs) is the ratio of the likelihood of one particular hypothesis to the likelihood of another. We categorise BFs based using the heuristic classification scheme for BF_10_ (Lee and Wagenmakers, 2013, p.105; adjusted from Jeffreys, 1961). Thus, for example, BF_10_ = 10-30 denotes strong evidence, BF_10_ = 3-10 moderate, and BF_10_ = 1-3 anecdotal evidence for H_1_, while BF_10_ = 1/3-1 denotes anecdotal, BF_10_ = 1/10-1/3 moderate, and BF_10_ = 1/30-1/10 strong evidence for H_0_. Outliers were identified for each correlation separately by bootstrapping the Mahalanobis distance (Schwarzkopf et al., 2012). To statistically compare correlations, Fisher’s z-transformation was applied to each correlation coefficient, resulting in normally distributed values *r’* with standard errors *s*_*r’*_. The null hypotheses (*r*^*’*^_1_-*r*^*’*^_2_=0) were tested in R(psych) (Revelle, 2015) using Student *t* test (Howell, 2011). Reported *p*-values are 2-tailed.

## 3 Results

### 3.1 MEG reveals expected ERD and ERS in the beta band

Clear movement-related changes in power were observed in all three frequency bands (β, slow-γ, mid-γ), characterized by different spectra-temporal-spatial properties. We observed a clear β ERD during movement and a β ERS after movement termination. In line with previous studies, the β ERD started before movement onset, plateaued between movement onset and offset, and terminated after movement offset (**Fig. 2a**, bottom left). The mean β ERD peak frequency was 19.33 Hz (range 15-26 Hz). There was moderate evidence for a lack of relationship between peak frequency and power (*r* = 0.011, BF_10_ = 0.22) across individuals. Again, consistent with prior observations, the β ERS started after movement offset and lasted for roughly 1 s (**Fig. 2a**, bottom right). The mean β ERS peak was 18.21 Hz (range 14-25 Hz). There was anecdotal evidence for a lack of relationship between peak frequency and power (*r* = -0.284, BF_10_ = 0.74).

**Fig. 2.**
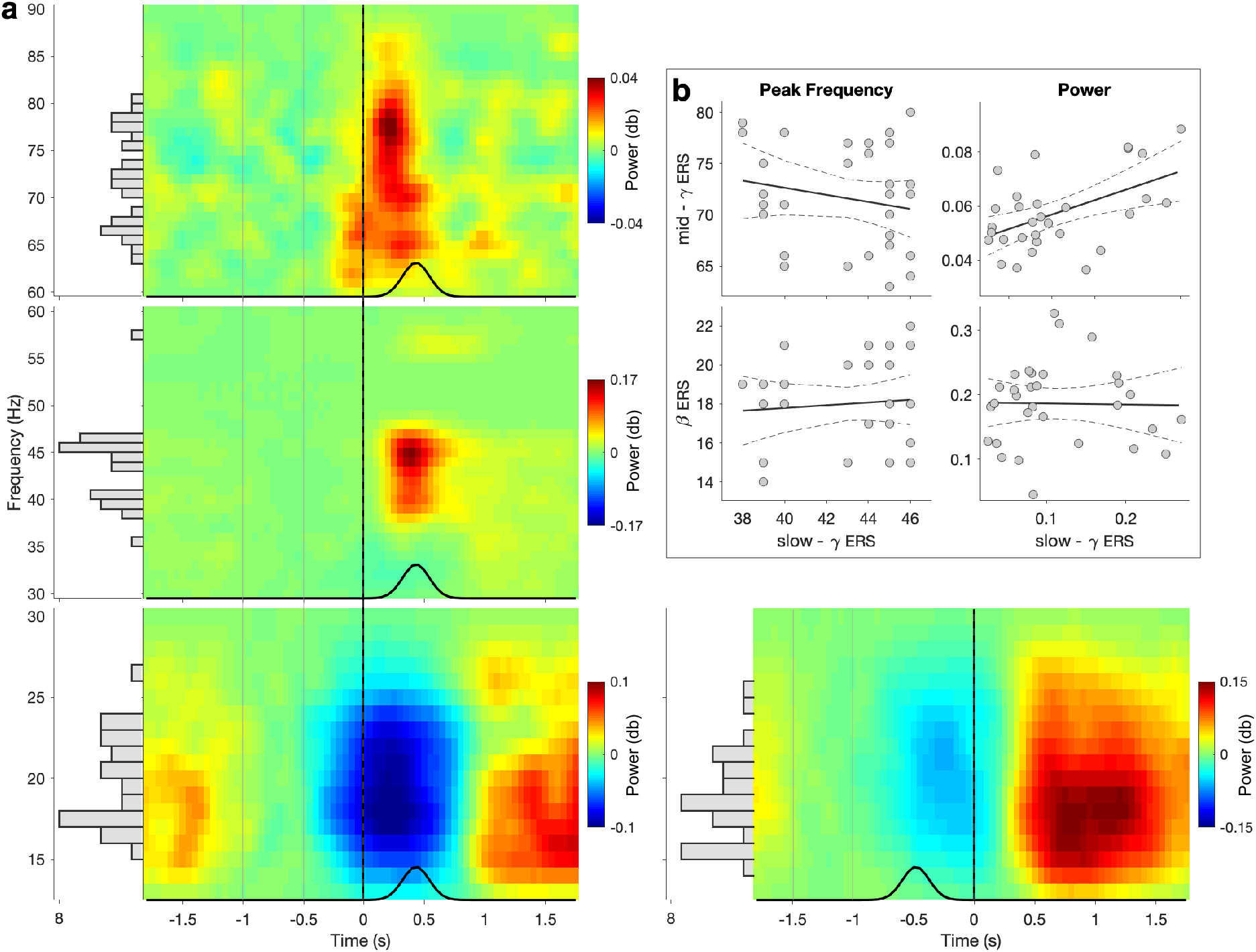
Temporal and spectral properties of movement-related responses. (a) Changes in power relative to baseline (−1 to -0.5 s relative to movement onset [-1.5 to -1 s relative to movement offset for β ERS], as indicated by the grey vertical lines). Data are locked to movement onset (movement offset for β ERS, as identified by EMG and highlighted by the black vertical line). Black line represents the distribution of movement offsets (movement onsets for β ERS) of trials included in the analysis. Side panel histograms illustrate the distribution of individual peak frequency (bin size = 1 Hz). (b) Correlations for peak frequency (left) and power (right) between slow-γ ERS and β ERS (bottom) as well as slow-γ ERS and mid-γ ERS (top). Dashed lines represent the 95% confidence intervals.

### 3.2 Two distinct patterns of movement-related γ activity

We then wanted to investigate movement-related activity in the γ bands. In the slow-γ band we observed a strong ERS, which started after movement onset, reached its peak at the time of movement offset, and decreased after movement offset, suggesting that the slow-γ ERS was temporally aligned with movement offset (**Fig. 2a**, center left). The mean slow-γ peak frequency was 43.06 Hz (range 35-57 Hz), and moderate evidence for a lack of relationship between peak frequency and power was found (*r* = -0.034, BF_10_ = 0.22). In the mid-γ band we also observed an ERS. This ERS started at movement onset, reached its peak between movement onset and offset, and terminated around movement offset (**Fig. 2a**, top left), therefore showing a temporal alignment with movement, unlike the slow-γ ERS. The mean mid-γ peak frequency was at 71.36 Hz (range 63-80 Hz), and there was anecdotal evidence for a lack of relationship between peak frequency and power (*r* = 0.249, BF_10_ = 0.55).

To our knowledge, while movement-related slow-γ has been reported previously (Crone et al., 1998; Szurhaj et al., 2006), its properties have not been fully characterised. We therefore sought to investigate whether this pattern of neural activity was distinct from the movement-related β ERS and mid-γ ERS. We performed four Bayesian pairwise correlations to test whether the peak frequency or power of the slow-γ ERS was related to these measures derived from the β ERS or mid-γ ERS (**Fig. 2b)**. We found moderate evidence for a lack of relationship between slow-γ ERS and β ERS peak frequency (*r* = -0.196, BF_10_ = 0.38) and anecdotal evidence for a lack of relationship between slow-γ ERS and mid-γ ERS peak frequency (*r* = 0.088, BF_10_ = 0.26). In terms of power, there was moderate evidence for a lack of a relationship between slow-γ ERS and β ERS (*r* = -0.019, BF_10_ = 0.22), but strong evidence for a relationship between slow-γ ERS and mid-γ ERS (*r* = 0.514, BF_10_ = 12.61).

Next, we examined the spatial properties of the movement-related slow-γ ERS compared with the β ERD, β ERS and mid-γ ERS. We considered the group-heatmaps of the peak virtual sensors and power maps. In line with previous findings, the group-heatmaps of the peak virtual sensors for β ERD and β ERS were relatively focal with the hotspot posterior and relatively central on the superior-inferior axis within the M1 (**Fig. 3a**). In contrast, the hotspot for the slow-γ was more superior and central on the anterior-posterior axis. Finally, the heatmap for the mid-γ was less focal, encompassing the hotspot of the β and the slow-γ frequency range.

**Fig. 3.**
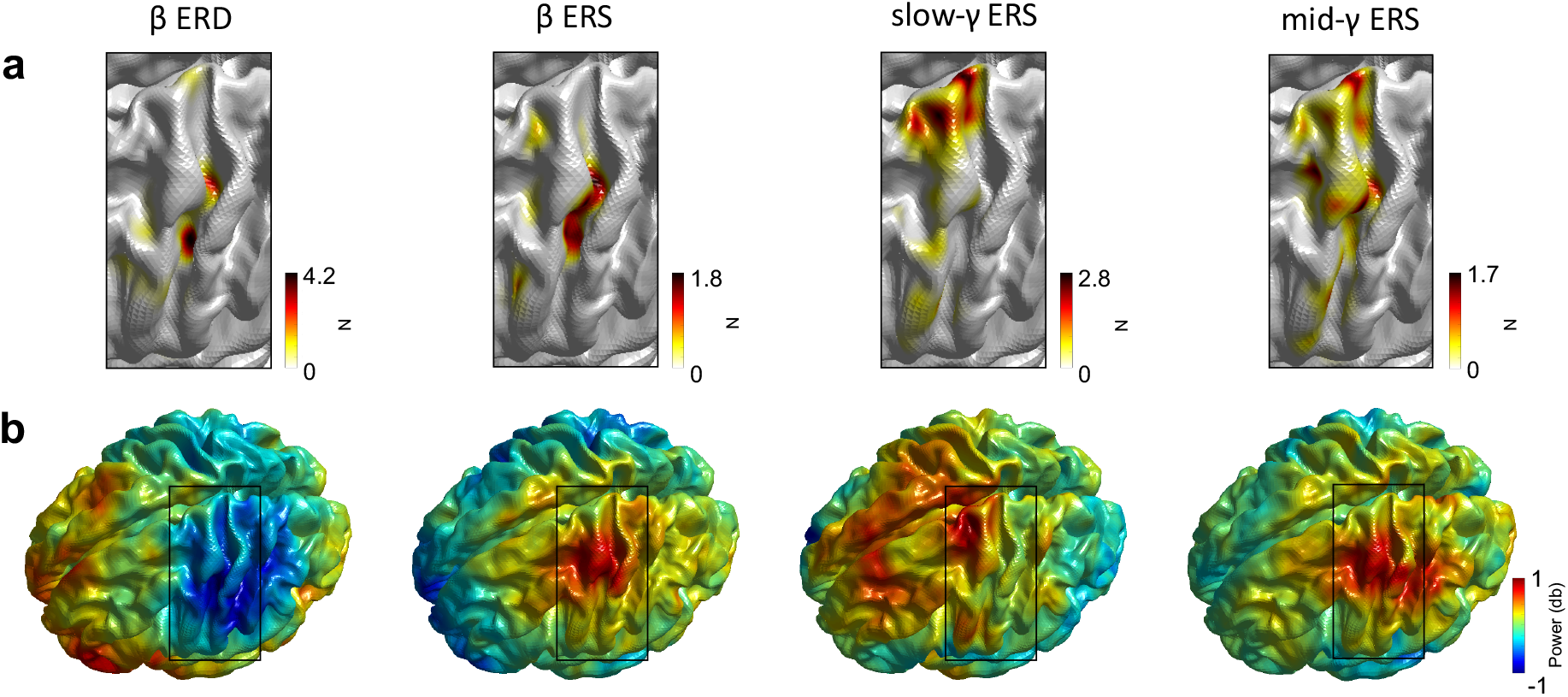
Spatial properties of movement-related responses. (**a**) Heatmap of the number of selected virtual sensors within M1. For visualisation data are interpolated. (**b**) Power maps, i.e. power averaged from movement onset to movement offset (movement offset to movement offset + 1 s) at each of the of the 3559 virtual sensors at the individuals’ peak frequency (i.e. frequency with strongest positive [negative for β ERD] power change from baseline within M1).

The spatial pattern of the power maps was qualitatively comparable for β ERD, β ERS and mid-γ with peaks in central on the superior-inferior axis and the anterior-posterior axis within M1 (**Fig. 3b**). In contrast, the spatial map of the slow-γ was qualitatively more anterior within the M1 and extended to frontal areas.

### 3.3 Slow-γ ERS peak frequency is related to individuals’ GABA-A activity during movement preparation

Next, we investigated the neurophysiological underpinnings of the movement-related activity we observed. In line with our hypothesis that movement-related activity in the γ band is related to movement-related GABA-A activity, we found strong evidence for a positive relationship between pre-movement SICI amplitude and slow-γ peak frequency (*r* = 0.677, BF_10_ = 18.13). There was moderate evidence for a lack of relationship between pre-movement SICI amplitude and peak frequency in other bands (β ERD: *r* = -0.010, BF_10_ = 0.29; β ERS: *r* = 0.061, BF_10_ = 0.29; mid-γ: *r* = 0.07, BF_10_ = 0.295, **Fig. 4**, left). The observed relationship between pre-movement SICI and slow-γ peak frequency was significantly different from the relationships observed in other bands (SICI and slow-γ peak frequency vs SICI and β ERD peak frequency: *z* = -2.26, *p* = 0.024; SICI and slow-γ peak frequency vs SICI and β ERS peak frequency: *z* = -2.10, *p* = 0.036; SICI and slow-γ peak frequency vs SICI and mid-γ peak frequency: *z* = -2.07, *p* = 0.039).

**Fig. 4.**
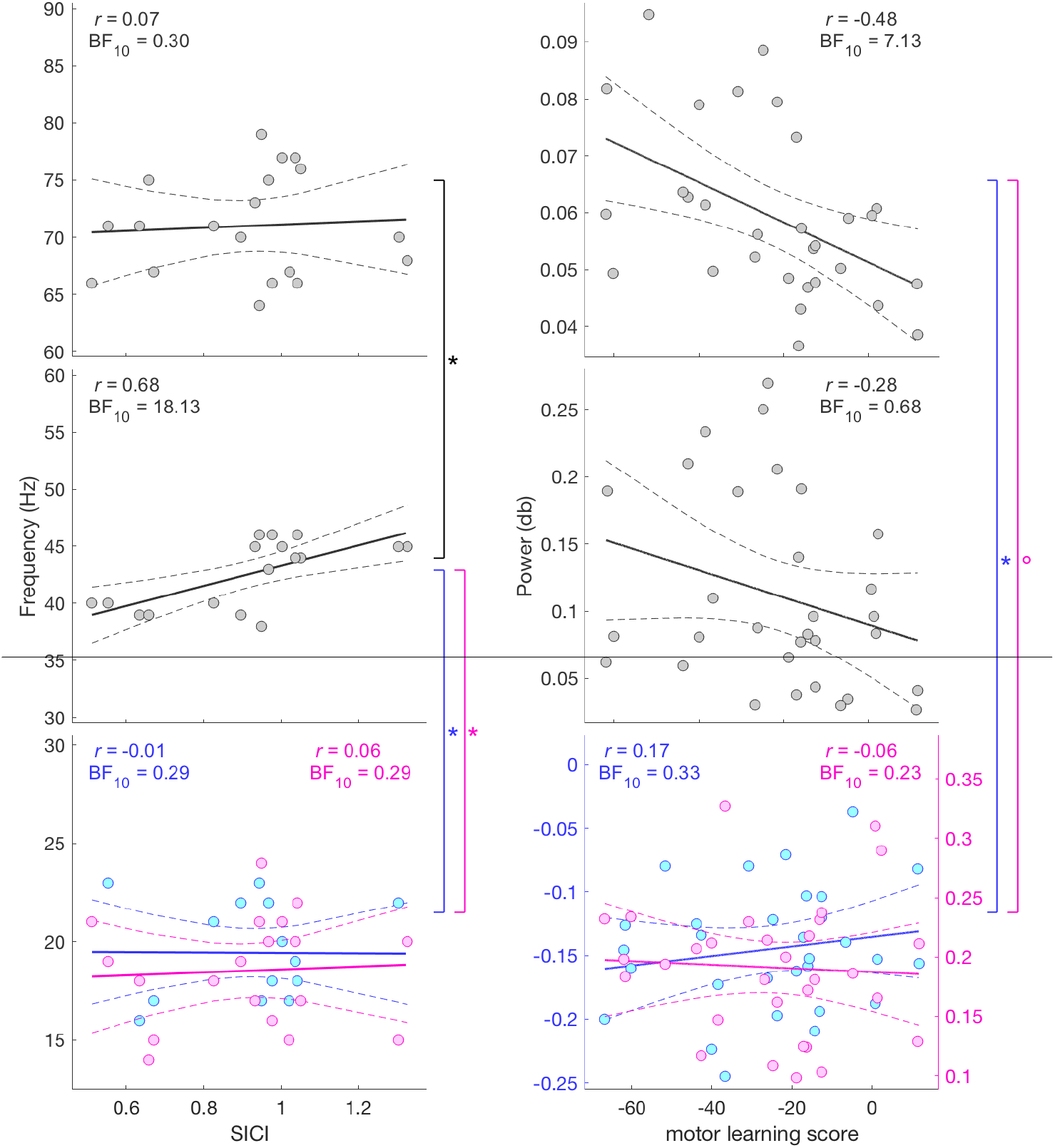
Relationship between MEG peak frequency and SICI amplitude (left) and MEG power and motor learning score (right) for β ERD (bottom blue), β ERD (bottom pink), slow-γ (middle) and mid-γ (top). Dashed lines represent the 95% confidence intervals. ° *p* < 0.1, * *p* < 0.5.

Having demonstrated a significant relationship between pre-movement SICI and slow-γ peak frequency, we then wished to explore the temporal specificity of this effect. Therefore, we investigated the relationship between pre-movement SICI and slow-γ peak frequency, for SICI_early_ and SICI_late_ separately, in post-hoc analyses. There was anecdotal evidence for a positive relationship between slow-γ peak frequency and GABA-A activity early in movement preparation (SICI_early_, *r* = 0.433, BF_10_ = 1.78) and strong evidence for a positive relationship between slow-γ peak frequency and GABA-A activity late in movement preparation (SICI_late_, *r* = 0.641, BF_10_ = 10.41). There was no significant different between these two correlations (*z* = -0.84, *p* = 0.401). There were no significant correlations between the peak frequency of any band and subsequent motor learning.

### 3.4 Mid-γ power correlates with subsequent motor learning

Finally, we wished to investigate the behavioural importance of these movement-related signals.

Firstly, it was important to determine whether participants were able to learn the task. As expected, RT decreased significantly across successive sequence blocks (*F*_(14,336)_ = 9.015; *p* < 0.001). In contrast, there was no significant difference in mean RT between the two random blocks (*t*_(32)_ = 0.885; *p* = 0.383), whereas there was a significant difference between block 14 (the final learning block) and block 15 (the second random block) (*t*_(28)_ = -6.899; *p* < 0.001), suggesting that improvements in RT occurred via learning of a specific sequence and not generic skill learning. There was also no significant difference between the RT from blocks 10-14, which were on the plateau of the learning curve (*F*_(4,108)_ =0.440; *p* = 0.780).

In line with our hypothesis, we found moderate evidence for a negative relationship between motor learning score and mid-γ power (*r* = -0.481, BF_10_ = 7.13, **Fig. 4** right), such that higher mid-γ power was related to greater motor learning. There was moderate evidence for a lack of relationship between motor learning score and power in other bands (β ERD: *r* = 0.166, BF_10_ = 0.33; β ERS: *r* = -0.056, BF_10_ = 0.23; slow-γ: *r* = -0.281, BF_10_ = 0.684). The observed relationship between motor learning score and mid-γ power was different from the relationship between motor learning score and β ERD power (*z* = -2.55, *p* = 0.011) and β ERS power (*z* = -1.72, *p* = 0.085). There were no significant relationships between the power in any band and SICI.

## 4 Discussion

This work aimed to examine the physiological basis and functional significance of movement-related M1 γ activity. We identified two distinct patterns of movement-related y activity in M1, characterised by different temporal-spectral-spatial properties. We went on to investigate the physiological correlates of these, and identified a correlation between M1 slow-γ peak frequency and pre-movement SICI amplitude in M1, such that individuals with a higher slow-γ peak frequency showed less GABA-A activity. Finally, in line with previous work we found that a higher M1 mid-γ power was related to better individual performance in a motor learning task.

### 4.1 Two distinct movement-related patterns of γ activity

Animal studies and human direct cortical recordings have suggested the presence of two distinct patterns of activity within the γ band in M1, but until now it has proved difficult to robustly separate them with transcranial approaches. By optimising our MEG analysis, i.e. separate beamformer for each sub-band and high-precision peak frequency/spatial location estimation, we have demonstrated the presence of two movement-related γ activity patterns within M1.

Given the paucity of previous transcranial studies focussing on slow-γ movement-related activity, we first examined whether the slow-γ activity seen here represented a distinct neural activity pattern, or was merely an extension of either post-movement β ERS or movement mid-γ ERS. We then reasoned that if the slow-γ ERS reflected activity within the same local microcircuits as either the post-movement β ERS or movement mid-γ ERS we would expect to observe systematic relationships on a subject-by-subject basis between slow-γ and β or mid-γ. We only found a relationship between slow-γ ERS power and mid-γ ERS power. The absence of other relationships add weight to the hypothesis that the slow-γ ERS is a distinct movement-related activity pattern but is in itself not conclusive. We therefore went on to investigate both the temporal and spatial domain of the slow-γ ERS, demonstrating that it is dissociable from both the post-movement β ERS and movement mid-γ ERS in both of these domains. Slow-γ ERS appeared to be temporally aligned with the movement offset, rather than movement onset, like the mid-γ ERS, or post-movement, like β ERS, and has a spatial distribution that is more frontal than either the mid-γ ERS, which is more localised to M1 or the beta-ERS.

Taken together, the data suggest that two distinct patterns of movement-related γ activity are seen in the human M1. The next question we aimed to address is the likely cellular basis of these two distinct γ activity patterns.

### 4.2 Slow-γ activity likely arises from superficial cortical layers

A commonly held hypothesis states that activity in the lower cortical layers is predominantly slower than that in the more superficial layers (i.e. frequency-layer gradient), reflecting the different functional roles of superficial and deep layers. This is supported by animal studies in primary sensory areas (e.g. (Roopun et al., 2006; Buffalo et al., 2011; Spaak et al., 2012; Haegens et al., 2015)). Moreover, human laminar MEG showed that visual α activity and sensorimotor β activity localise more to the white matter surface approximating infragranular origin than to the pial surface, while visual and sensorimotor γ activity localise more to the pial surface approximating supragranular origin than to the white matter surface (Bonaiuto et al., 2018). However, there is also evidence challenging the frequency-layer gradient by suggesting deeper cortical layers as origin for γ activity. For example, in the visual cortex of behaving mice, γ activity has been linked to parvalbumin (PV)-positive GABAergic interneurons (Chen et al., 2017), which are most densely populated in layer V (Fagiolini et al., 2004; Sohal et al., 2009). Further, auditory *in vitro* work revealed two distinct γ activities, 30-45 and 50-80 Hz, originating in layer II/III and layer IV, respectively (Ainsworth et al., 2011). The precise neural basis of γ activity in the primary M1, as opposed to primary sensory regions, has yet to be determined (Whittington et al., 2011). Translating the findings directly from studies performed in the sensory areas to M1 must be done with care, as the circuit organisation of M1 differs fundamentally from that of sensory areas, not least in that it is agranular, lacking a distinct layer IV (Shipp, 2005; Shipp et al., 2013).

To explore whether M1 movement-related β and/or γ activity arises from superficial cortical layers, we tested the relationship between β, slow-γ and mid-γ and SICI amplitude, a direct measure of local cortical GABA-A activity, quantified via TMS. TMS preferentially stimulates more superficial neurons, particularly at the intensities used here (Siebner et al., 2009). Further, computational modelling studies have demonstrated that TMS effects can be explained by activity within the canonical microcircuit, which includes layer II/III and layer V excitatory pyramidal cells, inhibitory interneurons, and cortico-cortical and thalamo-cortical inputs (Di Lazzaro and Ziemann, 2013). We demonstrated a specific relationship between local GABA-A activity and slow-γ activity, which was not observed for either the β or mid-γ activity. Importantly, different microcircuits within M1 have distinct patterns of GABA-A receptor morphology in terms of their α subunits. SICI has been demonstrated to primarily represent activity within cortical microcircuits involving interneurons that express GABA-A receptors with α-2 and α-3 subunits, rather than α-1 (Di Lazzaro et al., 2007). α-1 subunits are found at highest density in layer V of the healthy human M1, whereas α-2 subunits are more common in the superficial layers, and α-3 are fairly equally distributed across the cortical layers (Freund, 2003; Petri et al., 2003).

In summary, the data presented here suggest that movement-related slow-γ activity arises from neuronal circuits containing layer II/III interneurons. The functional role of movement-related slow-γ activity is less clear. Previous work has postulated that it may directly reflect motor output (Crone et al., 1998).

### 4.3 Mid-γ activity may reflect activity in learning-related microcircuits

In light of the significant correlation between slow-γ and SICI amplitude the absence of the same relationship for mid-γ could be interpreted in at least two ways. First, in human M1 slow-γ and mid-γ steam both from superficial layers, but from different populations or microcircuits, with the one underlying slow-γ, but not mid-γ, being GABAergic as measured using SICI. This would be in line with the frequency-layer gradient reported in sensory areas (Roopun et al., 2006; Buffalo et al., 2011; Spaak et al., 2012; Haegens et al., 2015; Bonaiuto et al., 2018).

Second, while slow-γ arises superficially mid-γ arises from deeper layers, such as layer V. This is in conformity with other animal work in sensory areas (Sohal et al., 2009; Ainsworth et al., 2011; Chen et al., 2017) and the supported by the functional role of M1 mid-γ. We demonstrated that the power of mid-γ activity elicited by a simple movement predicted the ability to learn a motor skill on a subject-by-subject basis. This is in line with previous work. For example, we have shown that an individual’s response to 75 Hz tACS relates to their ability to learn a subsequent motor skill (Nowak et al., 2017), and further, when amplitude modulated by an underlying theta pattern, improves motor learning in healthy adults (Akkad et al., 2019). The finding of a specific relationship between mid-γ activity and plasticity is consistent with data from animal recordings suggesting that microcircuits containing α-1 GABA-A synapses, predominantly found in the PV-rich layer V in M1, are a major neural substrate for plasticity, at least in the visual cortex (Fagiolini et al., 2004). Together, the origin of mid-γ activity is not completely understood, but mid-γ activity seems to play a role in motor learning.

### 4.4 Limitations

This study used non-invasive recordings to indirectly study changes in movement-related activity in the motor cortex. While this approach provides an unrivalled ability to understand activity in the healthy human system, it has inherent limitations in terms of the conclusions we can draw. Specifically, here, it was difficult to accurately quantify activity around 50 Hz due to power line noise. In addition, due to an artefact caused by the HPI coils at 53 and 54 Hz, we had to widen the standardly-employed line noise notch filter to 49-55 Hz. This had direct implications on our assessment of the peak frequency of slow-γ ERS.

### 4.5 Conclusions

The findings presented here allow us to create a theoretical framework for γ activity in the human M1, as follows: there are two patterns of movement-related γ activity in the human motor cortex (slow-γ and mid-γ), with differential temporal, spectral and spatial properties. The frequency of movement-related slow-γ activity is related to our neurophysiological measure of GABA-A activity, but does not play a direct role in motor plasticity *in vivo*. We conclude that slow-γ arises from alpha-2 and alpha-3 GABA-A microcircuits in layer II/III. Mid-γ activity predicts motor learning, but weather it originates layer II/III or V is not yet clear. This framework draws together findings from this paper and the literature, and provides a number of hypotheses that can be directly tested.

## Acknowledgements

We thank Miles Whittington for helpful discussions on the functional role of gamma activity.

## Data availability

Data are available from the first authors and the last author upon request.

## Author contributions

MN and CJS designed the study. MN, EH and CZ collected the data. CZ and MN analysed the data. AJQ and MWW provided assistance with data analysis. CJS supervised the project. CZ, MN and CJS wrote the manuscript and EH, AJQ and MWW edited the manuscript.

